# Male germline vulnerability to DNA damage causes sex-biased mutation

**DOI:** 10.1101/2025.08.20.671249

**Authors:** Danqi Qin, Laurence D. Hurst, Haoxuan Liu

## Abstract

In many vertebrates the male germline mutation rate is higher than that of females, so-called male mutation bias, the causes of which are central to numerous molecular evolution debates including the relationship between life history traits and the mutation rate and the cause of the generation time effect. The male bias was classically considered a consequence of the higher number of cell divisions in the male germline, but recently multiple lines of evidence find this to be an inadequate explanation. The revived damage-induced hypothesis posits that males are either more prone to germline damage or less likely to repair damage, leading to more mutations. The model predicts that, when males and females are treated with approximately equal levels of mutagenic stress, the male germline will accumulate more mutations. Here we provide the first test of the damage hypothesis by exposing male and female mice to mutagenic chemicals with both untreated and a non-mutagenic (PBS) controls, assaying *de novo* mutations via parent-offspring whole-genome sequencing. As predicted, in the instances where the mutagens caused a net increase to the mutation rate, it went up more in male germline. Comparison with established mutational signatures reveals differential involvement of the base excision repair (BER) pathway. Similarly, transcriptome analysis finds higher expression levels of DNA repair genes in ovary compared to testis for those genes in the BER pathway, a difference that becomes more extreme on administration of mutagens. As parental age, hence number of germline cell divisions, is controlled, we conclude that the male mutation bias is, at least in part, owing to differential damage repair with lower base excision repair in males playing a significant role.

## Main

Human *de novo* inherited mutations (DNMs) are generated more from fathers than from mothers (*1*). This phenomenon of male-biased germline mutation was first documented by Haldane through analysis of haemophilia inheritance (*1*), strongly supported by comparison of putatively neutral substitution rates between sex chromosomes and autosomes (*2–4*) and confirmed using whole-genome sequencing, revealing a male-to-female germline mutation rate ratio (•) of approximately 4 in humans (*5*). The bias is widespread across vertebrates, with • values ranging from 1.5 to 7.6 among species (*6*), this leading to the concept of male-driven evolution (*3*) and questioning the mutational deterministic model for the evolution of sex (*7*).

Historically, male-biased mutation was explained by the replication-driven hypothesis (*3*, *8*) , a related hypothesis postulated to explain between-tissue variation in cancer rates (*9*). The replication model for male-biased mutation posits that a greater accumulation of replication errors occurs in the male germline because it continues to divide throughout the reproductive lifespan, while female germline has many fewer divisions and ceases mitotic activity before birth, the same logic being given to explain the increase in the mutation rate with male age. Under the assumption that replication is the source of the male bias, the bias has been important in debates concerning, for example, the relationship between life history and mutation rate (*10–12*) and between generation time and the rate of molecular evolution (*3*, *13*) – the so-called generation time effect.

While this explanation has been the accepted model for many years (*14*), recently several lines of evidence have cast doubt on its validity. For example, mutation rates increase over time in mothers as well as fathers (*15*, *16*). Related, despite the very different profiles of replication in male and female germline, the male-to-female mutation ratio is around 3.5 at puberty and increases very little with parental age (*17–34*). Looking across mammals, the male female mutation ratio is around 2-4 no matter how long after birth the species reproduces, which varies from months to decades, with estimates of the number of germ cell divisions varying also over a much broader scale (*23*). For example, direct estimate of • in baboons find it to be nearly identical to that in humans, even though baboons have much shorter generation times and are expected to have reduced male-to-female germline division ratios (*35*). In birds, differences between taxa in the degree of male mutation bias when estimated from rates of evolution of Z and W chromosomes appear to reflect differences in the rate of mutation on the female-specific W chromosome, an effect that cannot be attributed to the number of replications in male germline (*12*). The lack of evidence for replication-dependent mutation accords with evidence that post-mitotic neurons accumulate mutations at approximately the same rate as somatic cell types that are still replicating (*36*). Likewise, cell division numbers and mutational burden are uncoupled in mammalian colonic crypts (*37*) and as many as 90% of yeast’s mutations are not replicative in origin (*38*).

Given these deficiencies, focus has shifted to an alternative model, the differential damage hypothesis. As initially proposed by Haldane and recently revived (*1*, *33*, *39*), male-biased germline mutations could be explained by differential degrees of unrepaired DNA damage. Under this model, the stable • observed across age groups in humans can be reconciled if most germline mutations are damage-induced, that such damage accumulates as a function of time in both males and females, and the male’s ability to repair damage is consistently lower than in females. That both males (*40*) and females (*41*) have reduced germline damage repair as they age is consistent with such age effects.

In principle there could be two different, not mutually exclusive, forms of the damage hypothesis. One postulates that for any given dose of mutagen the male germline is more exposed to the mutagen but that investment into repair is no different to that in females. If a certain proportion of damage goes unrepaired, males would then have a higher mutation rate owing to the higher rate of damage. The alternative is that exposure is comparable in males and female germ line, but males invest less into repair. Males and females then have similar levels of damage but different degrees to which damage goes unrepaired. Supporting the former exposure model, protamine packaging in sperm may specifically lead to oxidative damage requiring repair in the early embryo (*42*). Consistent with the second model, sex-specific differences in mutation spectra, which reflect underlying mutational processes, may indicate differential damage repair efficiencies (*15*, *43*). The former may be considered a differential exposure model, the second a differential repair model, collectively, the damage hypothesis.

The damage-hypothesis has yet to be experimentally tested (in either subform or more generally). A central prediction is that if equal doses of a damage inducing agent are given to males and females, the male germline mutation rate should increase more owing to weaker protection (i.e. greater germline exposure per unit dose) and/or weaker damage repair (i.e. damage occurs at equal rates but is less likely to be repaired in males). Here we present the first test of this prediction, using a single strain of laboratory mice (to prevent strain specific differences in mutation rate conflating analysis) with whole genome parent-offspring sequencing employed to detect mutations. We find evidence supporting the damage-induced hypothesis, specifically in the differential repair mode, with males exhibiting significantly more raised mutation rates. Characterization of the signature of the extra mutations suggests involvement of base excision repair pathways, this supported by transcriptomic analysis.

### Experimental design

To detect germline mutations, mutagen-treated mice were paired with untreated mice to produce offspring, and parent-offspring trio sequencing was performed to identify *de novo* mutations in the germline (**Fig. 1A**). DNMs identified using this approach arise from two distinct origins: naturally occurring (black stars in **Fig. 1A**) and mutagen-induced (empty stars in **Fig. 1A**). Based on control experiments that determined the baseline frequency of naturally occurring DNMs, the number of mutagen-induced DNMs can be estimated by subtracting naturally-occurring DNMs from the total DNMs observed in the mutagen-treated trios (**Fig. 1A**).

**Figure 1.**
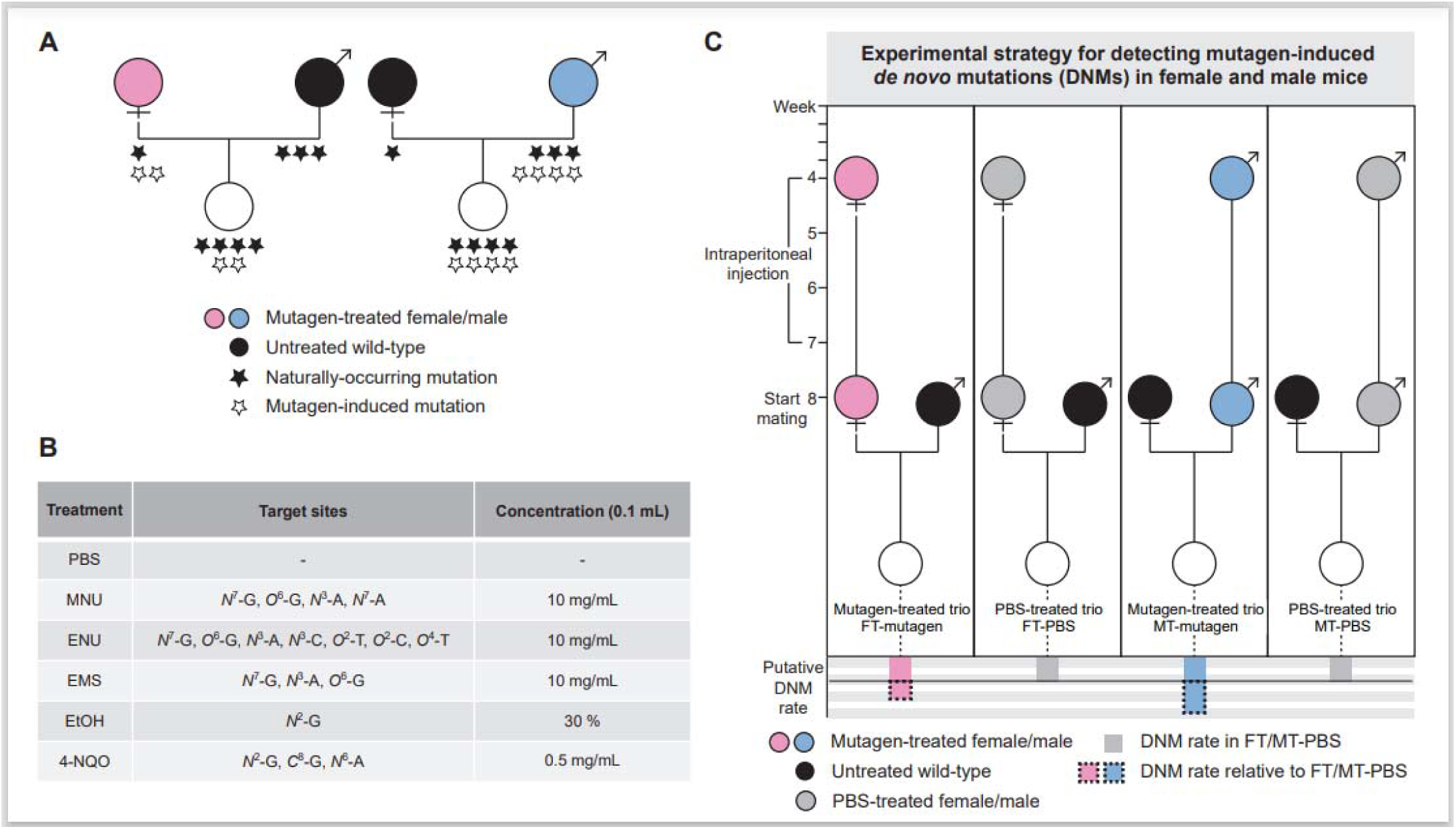
Experimental Design. (**A**) Schematic illustration of the hypothesis and resulting inference. A treated individual mates with an untreated individual to produce offspring. In the two trios (i.e., one offspring and its mother and father), parents of the same sex and age contribute comparable numbers of naturally-occurring germline mutations. Upon mutagen exposure, the male parent accumulates more mutagen-induced mutations in the germline (empty stars under the blue circle) than the female parent (empty stars under the pink circle). Mutations identified in the offspring consist of both naturally-occurring and mutagen-induced germline mutations, and the numbers differ between the two trios. (**B**) Table of treatments used in the experiment. Phosphate buffered saline (PBS) was the solvent for five mutagenic chemicals: *N*-Methyl-*N*-Nitrosourea (MNU), *N*-Ethyl-*N*-Nitrosourea (ENU), Ethyl Methanesulfonate (EMS), Ethanol (EtOH) and 4-Nitroquinoline-1-Oxide (4-NQO), all of which cause DNA damage by introducing structural alterations at specific nucleotide sites. The chemicals were delivered via intraperitoneal injection of 0.1 mL at its designated concentration. (**C**) Schematic illustration of the experimental strategy. Beginning at four weeks of age, with three-week intraperitoneal exposure followed by one-week recovery, each treated mice was mated with an age-matched untreated mice to produce the following four types of trios: females treated with mutagens (FT-Mutagen), females treated with PBS (FT-PBS), males treated with mutagens (MT-Mutagen), and males treated with PBS (MT-PBS). The DNM rate relative to FT/MT-PBS is defined as the DNM rate in each mutagen-treated trio minus that in the corresponding PBS-treated trio of the same sex.

As it is not expected to induce germ line mutations phosphate buffered saline (PBS) served as the solvent control, providing a baseline. We employed five extensively characterized, hence commonly used, mutagens. All are carcinogenic chemical agents that induce DNA damage by modifying different structural sites on the nucleotides (**Fig 1B**). Among these chemicals, *N*-Methyl-*N*-Nitrosourea (MNU), *N*-Ethyl-*N*-Nitrosourea (ENU), and Ethyl Methanesulfonate (EMS) are alkylating agents that react directly with DNA, transferring alkyl groups (primarily methyl or ethyl) to nitrogen or oxygen atoms in deoxyribonucleotides (*44–47*). In contrast, ethanol and 4-Nitroquinoline-1-Oxide (4-NQO) induce DNA damage through their metabolic products, which form covalent DNA adducts (*48–51*).

Equal doses of PBS or each of the five mutagens were administered to female and male C57BL/6J mice via intraperitoneal injection, starting at four weeks of age (**Fig. 1C**). As males are slightly heavier than females, the dosage per unit body mass will be slightly lower for males than females, rendering any increase in male mutation rate a conservative result. Treatments continued for three weeks, administered five times per week. Upon reaching eight weeks of age, each treated mouse was individually paired with an age-matched untreated mouse to produce offspring. Following reproduction, genomic DNA from both parents and offspring underwent whole-genome sequencing. By comparing the number of DNMs observed in offspring from trios with females treated by mutagen (FT-mutagen, FT denotes female-treated trios) to those from females treated by PBS (FT-PBS), we quantified the mutagenic effects specifically in female germline (the dashed box in **Fig. 1C**). Likewise, mutagenic effects in male germline were assessed by comparing offspring from trios with males treated by mutagen (MT-mutagen, MT denotes male-treated trios) to those from males treated by PBS (MT-PBS) (**Fig. 1C**).

### Male germline is more sensitive to mutagens

In each of the six treatment groups (PBS and five mutagens), five female and five male mice were included. In total, 51 out of the 60 treated mice successfully produced offspring when crossed with untreated mice. Both parents and at least one offspring from each of the 51 mating pairs underwent deep next-generation sequencing with an average depth of 39.6× (**table S1**). DNMs were identified by screening variants present in the offspring but absent in both parents (**table S2**) (see **Methods** for details). Seven DNMs were randomly selected for PCR verification, with 100% consistency observed with the next-generation sequencing results.

In the control group, the single nucleotide variant (SNV) rate for the PBS-treated mice was estimated at 4.2×10^-9^ (95% CI: 3.6×10^-9^ – 4.9×10^-9^) per site, consistent with previous estimates for untreated mice (3.9×10^-9^) (*52*), indicating that PBS treatment is not mutagenic. Furthermore, the SNV rates between the FT-PBS and MT-PBS were highly similar – 4.3×10^-9^ (95% CI: 3.4×10^-9^ – 5.3×10^-9^) vs. 4.1×10^-9^ (95% CI: 3.3×10^-9^ – 5.0×10^-9^) – consistent with expectations for naturally-occurring mutations in trios under the non-mutagenic treatment. This similarity reinforces the reliability of the PBS control as a stable baseline for evaluating mutagen-induced mutations across sexes.

In the mutagen-treated groups, we initially sequenced seven to nine trios under each treatment and found that the SNV rate was significantly higher in the male treatment (MT) under MNU and ENU treatments (Poisson tests, *P* = 0.0038 and 0.023 after Benjamini-Hochberg correction), while no significant differences were observed between FT and MT under EMS, EtOH, or 4-NQO treatments (**fig. S1**). The lack of noticeable sex-specific differences with these treatments may reflect that these mutagens did not exert sufficiently strong mutagenic effects, as the SNV rates under these treatments do not differ significantly from that in the control group (Poisson test, all *P* > 0.1; **fig. S2**). This lack of difference could be attributed to factors such as limitations in the dosing and adduct stability (*50*), as well as the possibility that male and female germlines may have similar repair abilities for certain lesions, since sex differences need not exist across all lesion types.

To further corroborate the incidences where a male bias is seen, and to collect more data for analysis of mutation spectrum, an additional 32 offspring from MNU-treated trios and 8 offspring from ENU-treated trios were randomly selected for sequencing. The resulting SNV rates were consistent with our previous results (**Fig. 2A**). Specifically, the SNV rate in MT-MNU averaged 6.6×10^-9^ (95% CI: 6.0×10^-9^ – 7.2×10^-9^) per site, significantly higher than the average of 4.0×10^-9^ (95% CI: 3.6×10^-9^ – 4.4×10^-9^) in FT-MNU (Poisson test, *P* < 0.001). The strongest mutagenic effect was observed with ENU treatment, where the SNV rate in MT-ENU reached an average of 2.6×10^-8^ (95% CI: 2.4×10^-8^ – 2.8×10^-8^) per site, approximately seven times higher than the control group, significantly higher than the average of 1.6×10^-8^ (95% CI: 1.5×10^-8^ – 1.7×10^-8^) in FT-ENU (Poisson test, *P* < 0.001).

**Figure 2.**
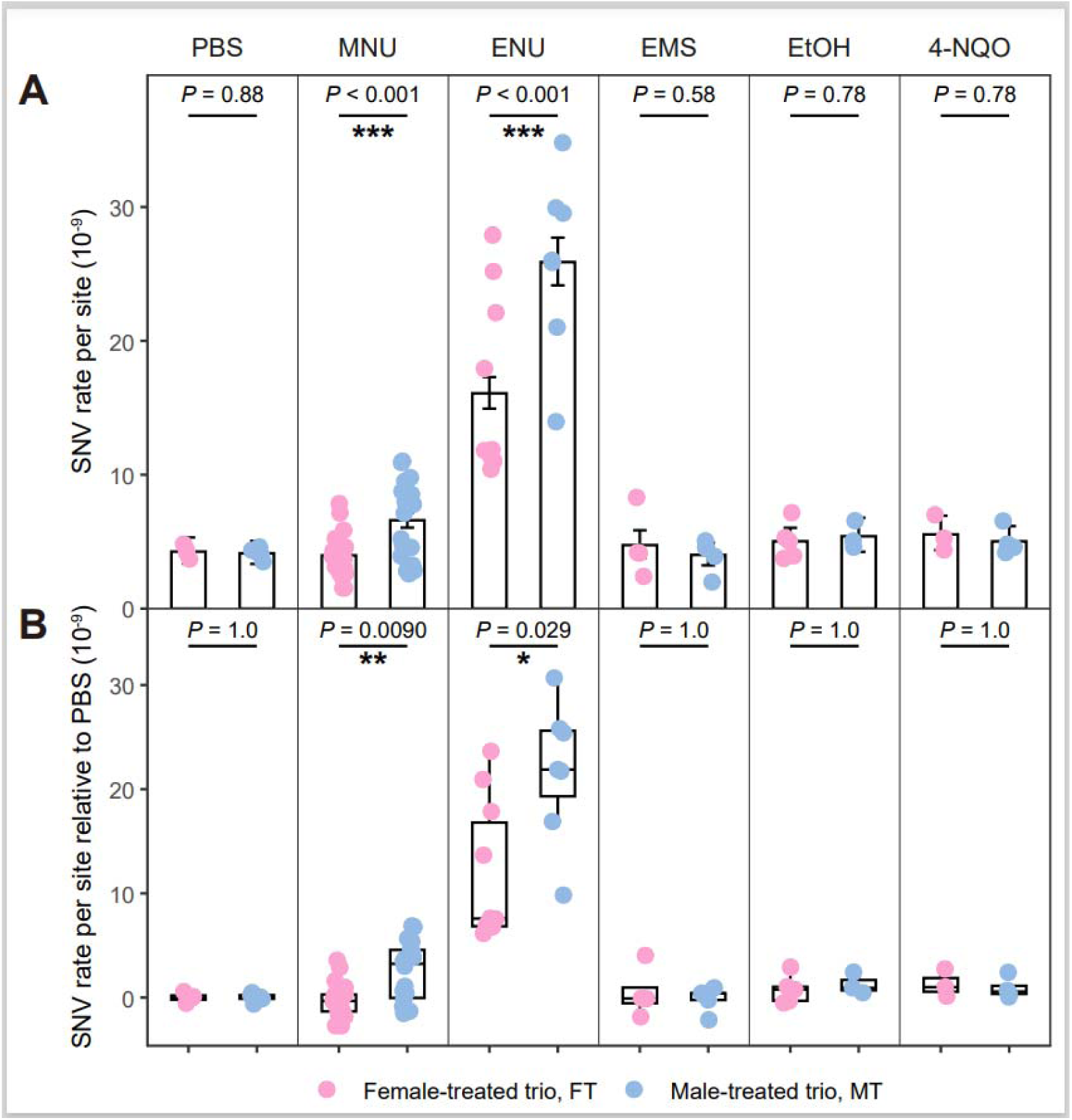
Comparison of germline SNV rates between FT and MT under each treatment. (**A**) Sex-specific germline SNV rates under six treatments. Histograms represent the average SNV rates, with error bars indicating the 95% confidence intervals predicted by Poisson distribution. *P* values are obtained using the Poisson test, and ‘***’ denotes *P* < 0.001. (**B**) Sex-specific germline SNV rates after subtracting the mean SNV rate of the corresponding sex’s PBS control group. Boxes represent the interquartile range (IQR), with the central line indicating the median and whiskers extending to 1.5×IQR. Each point represents an offspring sample, jittered slightly to avoid overlap. *P* values are obtained using Wilcoxon rank-sum test, ‘*’ denotes *P* < 0.05, and ‘**’ denotes *P* < 0.01.

Sex parity remained consistent after accounting for the background SNV rate in the control group. After subtracting the SNV rate of the PBS control from those under mutagen treatments, MT still exhibited significantly higher SNV rates than FT under MNU and ENU treatments (Wilcoxon rank-sum tests, *P* = 0.0090 and 0.029 after Benjamini-Hochberg correction; **Fig. 2B**). Collectively, these results provide evidence that comparable levels of mutagenic stress induces more germline mutations in males than in females, supporting the damage-induced hypothesis.

### MNU- and ENU-treatment induced sex-specific mutation spectra

A specific mutagenic mechanism is often reflected in a characteristic mutation spectrum. To further investigate the mutational patterns induced by different mutagens and their potential sex-specific differences, we analyzed the relative frequencies of six types of SNVs across treatment groups. Within FT, a significant deviation from the mutation spectrum of the PBS control was observed only in FT-ENU (6×2 *•*² test, *P* < 0.001 after Benjamini-Hochberg correction; FT in **Fig. 3A**). In contrast, in MT, both MNU and ENU treatments showed significant differences from PBS (6×2 *•*² tests, all *P* < 0.001 after Benjamini-Hochberg correction; MT in **Fig. 3A**). Notably, under MNU treatment, FT exhibited no significant differences in either SNV rate or spectrum compared to FT-PBS, while MT showed significant differences in both SNV rate and spectrum compared to MT-PBS. This suggests that MNU treatment induces mutations specifically in the male germline, but not in the female germline, providing direct evidence of sex-specific mutagenic outcomes under equivalent treatment. Moreover, when comparing FT and MT under the same mutagenic exposure, MNU and ENU both exhibited significant between-sex differences (6×2 *•*² tests, all *P* < 0.001 after Benjamini-Hochberg correction), further indicating sex-specific responses to mutagenic stress.

**Figure 3.**
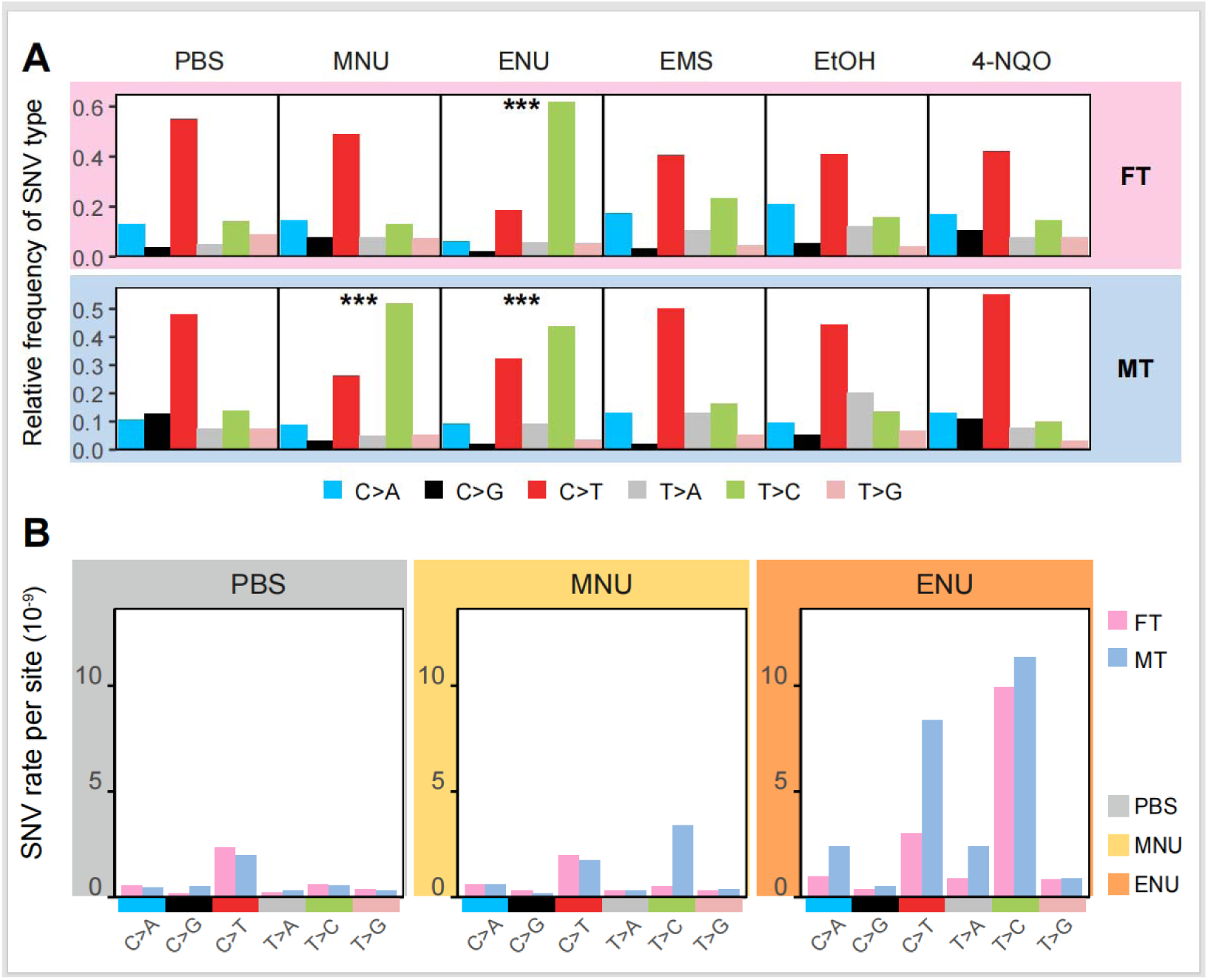
Germline mutation spectra under each treatment. (**A**) Mutation spectra under six treatments. Each facet in FT (female-treated trios) and MT (male-treated trios) rows shows the relative frequencies of six SNV types aggregated across from all samples under the respective treatment. ‘***’ denotes a Benjamini-Hochberg corrected *P* value < 0.001 compared to the sex-matched PBS control (*•*² test). (**B**) Rates of each SNV type per site per sample under three treatments: PBS (grey), MNU (yellow) and ENU (orange). Pink and blue represent female-treated and male-treated trios, respectively.

Nevertheless, the mutational spectrum reflects only the relative proportions of SNV types and does not capture the absolute number of mutations induced by specific mutagens. We thus examined the absolute rate of each SNV type across PBS, MNU, and ENU treatments. Compared to PBS, MNU treatment induced only one type of SNV (T>C) in MT, whereas ENU treatment led to an overall increase in most SNV types in both FT and MT, with effects consistently stronger in the MT (**Fig. 3B**). These findings indicate that mutagen-induced lesions differ between MNU and ENU treatments, and that the mutagenic effects of both treatments are more pronounced in males, as evidenced by increased SNV rates and altered spectra, suggesting greater vulnerability to DNA damage in the male germline.

### Mutation signatures associated with DNA repair pathway deficiency are enriched in the male germline

Building on the observed sex bias in the six-type mutation spectra under MNU and ENU treatments, we next examined more detailed mutation spectra, which classify SNVs by substitution type and the flanking nucleotides into 96 types of single base substitutions (SBS). This high-resolution approach could offer further insights into the underlying mechanisms of mutagenesis, since distinct patterns (termed signatures) in the 96-type mutational spectra have been characterized in cancer genomics. To date, over 80 different SBS signatures associated with various mutagenic processes have been identified (*53*, *54*), serving as molecular fingerprints for deciphering the origin of mutations. While analysis of the mutational spectra may thus be helpful in determining possible underlying causes of any male-female difference in our experiments and in nature, we don’t expect that the precise signature need be the same as that seen in nature as the mutagens in nature will no doubt be more heterogeneous.

We began by comparing the 96-type mutation spectra in MNU and ENU treatments with that in PBS. As observed with the six-type mutation spectra, we found a decrease in C>T mutations and an increase in T>C mutations in MT-MNU, FT-ENU, and MT-ENU (**Fig. 4A**). To explore the mechanisms behind these changes, we analyzed the exposure of known SBS signatures in mutation spectra under PBS, MNU, and ENU treatments.

**Figure 4.**
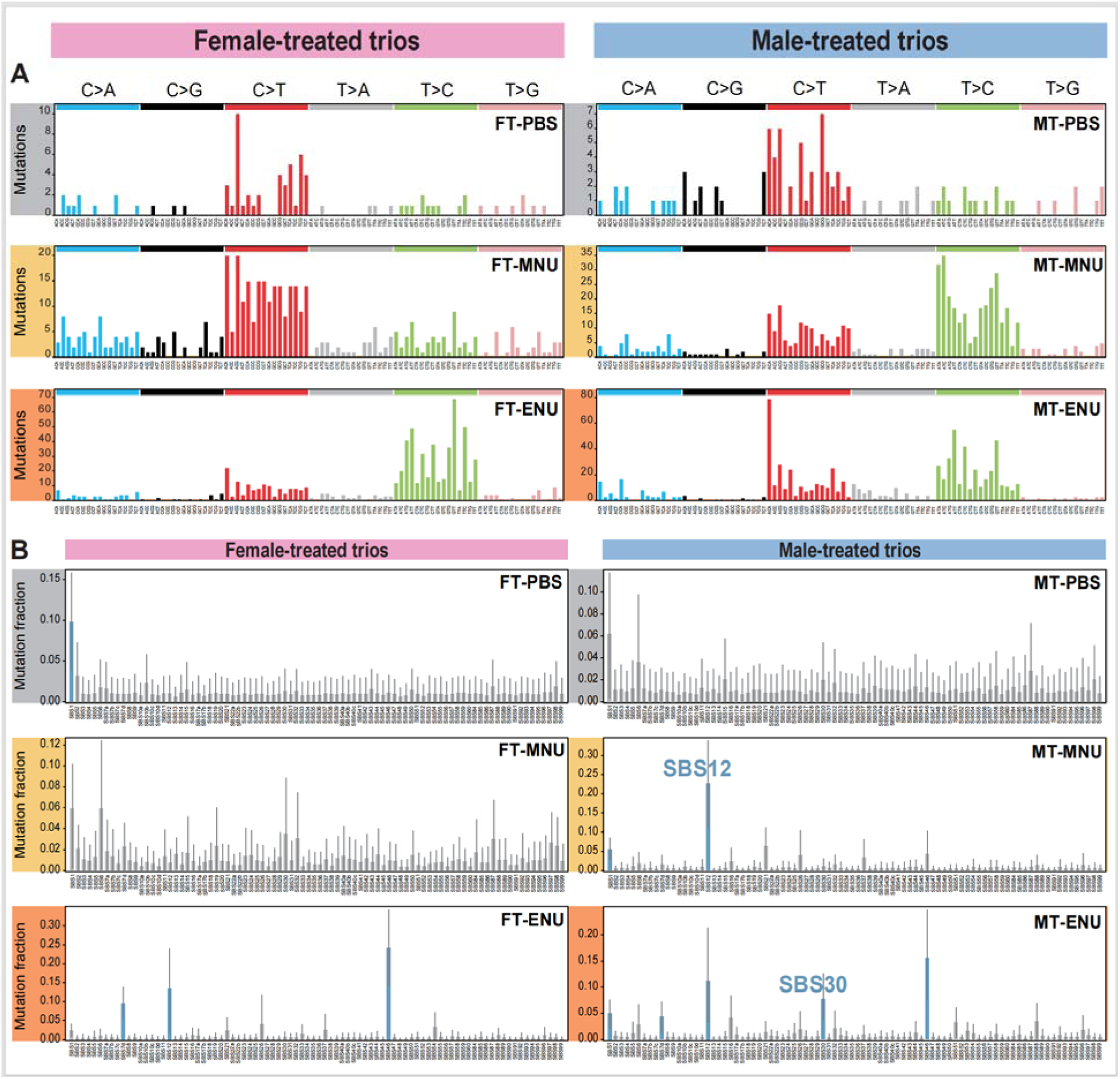
SBS mutational spectra and signature attribution for germline mutations under PBS, MNU, and ENU treatments. (**A**) SBS mutational spectra of female-treated and male-treated trios under three treatments. The *x*-axis shows 96 SBS types in their trinucleotide context, colored by base substitution type. The *y*-axis indicates the number of each mutation type. (**B**) SBS signatures attributed to the mutational spectra of female-treated and male-treated trios under each treatment. The *x*-axis shows 86 COSMIC mutational signatures. The *y*-axis indicates the level of exposure to each signature. Bars are colored blue for signatures with sufficiently non-zero estimated exposure and grey otherwise. Vertical lines represent the 90% highest posterior density (HPD) intervals, reflecting the uncertainty in the exposure estimates.

In the PBS control, compared with other signatures, we observed an increased contribution of SBS1 to the germline mutations in both FT and MT, with a more pronounced trend in FT (**Fig. 4B**). SBS1 is a common signature caused by the spontaneous deamination of 5-methylcytosine (*55*). Under MNU treatment, the contribution of SBS12, which is primarily characterized by T>C mutations, was substantially higher in MT but not in FT. Specifically, SBS12 accounted for over 20% of all mutations in MT-MNU, compared to only ∼1% in FT-MNU (**Fig. 4B**). This discrepancy suggests that a specific DNA repair pathway responsible for resolving SBS12-like mutations is deficient in males but remains functional in females. The molecular mechanism driving SBS12, however, is still unknown. Similarly, under ENU treatment, SBS30, which is primarily characterized by C>T mutations, contributed up to 10% of all mutations in MT but showed no discernible contribution in FT (**Fig. 4B**). SBS30 is caused by defects in base excision repair (BER), specifically with defects in NTHL1, suggesting that the BER pathway is more efficient in females than in males (and for this reason provide below focused analysis of NTHL1). The unique increase in mutations associated with SBS12 and SBS30 in males, but not in females, suggests that corresponding DNA repair pathways in the male germline are less efficient than in the female germline.

### Lower expression levels and weaker transcriptional activation of DNA repair genes in the male germline

The male-specific enrichment of SBS12 and SBS30 highlights the potential differences in DNA damage repair pathways between male and female germ cells. Since male and female mice share the same genetic background, this disparity may be owing to differences in the expression of DNA repair genes. To investigate this, we collected ovary and testis tissues from untreated mice and from mice treated with PBS, MNU, and ENU, and performed transcriptome sequencing. Our focus was on 214 genes known to be involved in DNA repair, spanning 16 categories of genes associated with DNA repair, including BER, nucleotide excision repair (NER), mismatch repair (MMR), homologous repair (HR), etc. (*56*) (**table S3**).

We initially compared the average expression levels of 214 genes between ovary and testis but found no significant difference under any condition (Paired-*t* test, all *P* > 0.17 after Benjamini-Hochberg correction). This lack of significance likely stems from substantial between-gene variation in expression levels, with the largest differences exceeding 1000-fold. To identify sex-biased gene expression, we then identified genes that were significantly differentially expressed between ovary and testis under each treatment (*t*-test, *P* < 0.05 after Benjamini-Hochberg correction; **table S4**), based on normalized expression levels measured as transcripts per million (TPM). Among the 214 DNA damage repair-related genes analyzed in untreated mice, 166 showed a significant sex bias with more at significantly higher levels in the ovary than in the testis (92 vs. 74), but this skew is not significant (Binomial test, *P* = 0.19) (NTHL1 is different in the two being much more lowly expressed in testis, *t*-test, *P* = 0.00046 after Benjamini-Hochberg correction). The difference between oocyte and testes in repair gene expression is, however, significant in control PBS-treated mice (112:66, Binomial test, *P* < 0.001). In mice treated with MNU and ENU, the sex-based disparity was even more pronounced, with ratios of 130:66 under MNU treatment (Binomial test, *P* < 0.001) and 131:65 under ENU treatment (Binomial test, *P* < 0.001) (**fig. S3**).

We can also consider the degree to which genes increase their expression on mutagen exposure. To evaluate the responsiveness of DNA repair genes under mutagenic exposure, we quantified the differential expression between treated and untreated conditions. As anticipated, mutagen treatments (MNU and ENU) induced significantly greater expression changes compared to PBS control (*t*-test, *P* < 0.001; **fig. S4**). Importantly, we observed a sex-specific transcriptional response where most DNA repair genes were upregulated in the ovary but downregulated in the testis following mutagen exposure (*t*-test, *P* < 0.001; **fig. S4**), highlighting a pronounced sex-based difference in transcriptional sensitivity to mutagenic stress. NTHL1 shows increased male female difference after mutagen exposure: the ratio of effect size of sex between PBS and untreated group is 1.4, but increased to 3.1 between MNU and untreated and 2.2 between ENU and untreated (**table S5**). Overall, the relatively lower expression and weaker transcriptional responsiveness of DNA repair genes in testes likely compromised functions related to DNA damage repair, contributing to an increased mutation rate in male germ cells.

Mechanistically, the damage induced by alkylating agents, such as MNU and ENU, is primarily repaired through the BER and NER pathways (*45*, *47*, *57*, *58*). Therefore, we focused on the 23 genes involved in the BER pathway and the 31 genes involved in the NER pathway (**table S3**). The greatest sex-based disparity was observed in the BER pathway. Of the 23 genes in the BER pathway, 15 were expressed at significantly higher levels in the ovary of untreated mice (Binomial test, P = 0.0075; **Fig. 5A**), and between 17 and 20 were expressed at higher levels in the ovary of PBS-, MNU-, and ENU-treated mice (Binomial tests, all P < 0.003; **Fig. 5A**). To evaluate the magnitude of between-sex transcriptional differences across treatment groups, we calculated Cohen’s d to measure effect size between ovary and testis in the untreated group, which served as our baseline control. We then determined the ratio of this effect size for each treated group relative to the untreated group (Cohen’s d treated / Cohen’s d untreated). Our findings indicate that sex-related transcriptional differences become more pronounced in the treated groups. Specifically, of the 17 BER genes that showed higher expression in untreated ovaries (Cohen’s d untreated > 0), the effect size increased for 16 genes in the PBS group, and for all 17 genes in the MNU and ENU groups (**Fig. 5B** and **table S5**). Moreover, the ratios of effect sizes for these 17 genes were significantly higher in the MNU and ENU groups compared to the PBS group (*t*-test based on log2-transformed values, both *P* < 0.001). This marked difference in gene expression likely contributes to the increased frequency of SBS30-associated mutations in MT-ENU. Notably, the expression level of NTHL1, the gene identified to cause SBS30 (*59*), in testis is only one-eighth of that in the ovary upon ENU exposure.

**Figure 5.**
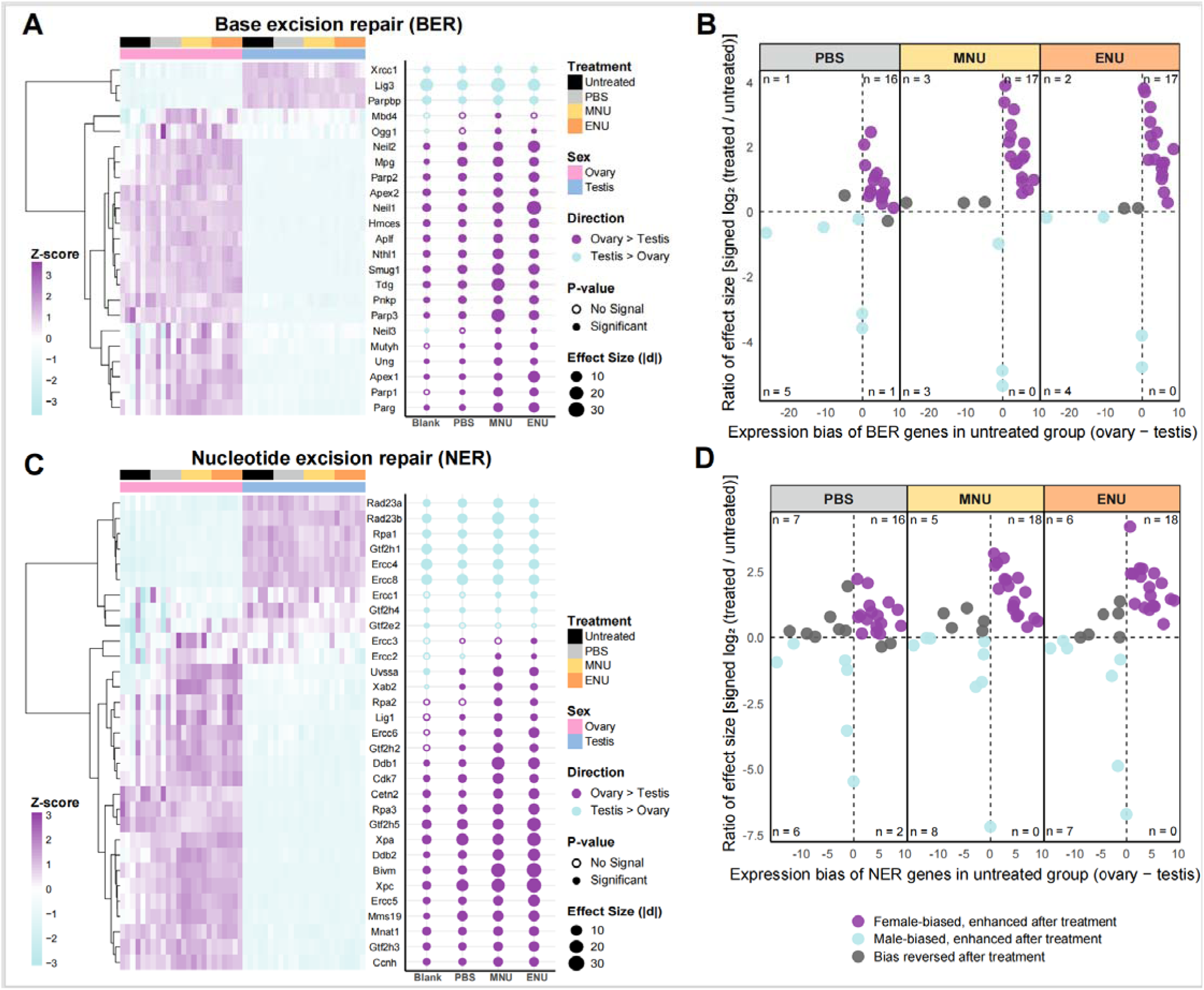
Expression patterns and sex-biased responses of DNA repair genes in the ovaries and testes. (**A**) Left: Heatmap showing BER gene expression in ovaries and testes under four conditions. Right: Dot plot illustrating effect size (Cohen’s d) for differences between ovaries and testes. (**B**) Scatter plots showing treatment-induced changes in sex-biased expression of BER genes. The x-axis shows effect size (Cohen’s d) between ovaries and testes in untreated mice. The y-axis shows the signed log_2_ ratio of effect sizes between treated and untreated groups, preserving the direction. Each point represents a gene. (**C**) Similar to (A), showing NER gene expression. For each heatmap, hierarchical clustering was performed using Euclidean distance and complete linkage. Rows and columns represent genes and samples, respectively, and color indicates standardized expression levels (Z-score). Z-scores are calculated by scaling TPM values by row. For each dot plot, each point represents a comparison of gene expression between ovaries and testes under a given treatment. (**D**) Scatter plots showing treatment-induced changes in sex-biased expression of NER genes. The x-axis shows effect size (Cohen’s d) between ovaries and testes in untreated mice. The y-axis shows the signed log_2_ ratio of effect sizes between treated and untreated groups, preserving the direction. Each point represents a gene.

For 31 genes associated with NER, the sex-based disparity in expression levels was not evident in untreated mice (Ovary higher: testis higher = 14:9; Binomial test, *P* = 0.40; **Fig. 5C**), while more genes were significantly highly expressed in the ovary of PBS-, MNU-, and ENU-treated mice (19:7 in PBS, 21:8 in MNU, and 22:8 in ENU; Binomial tests, all *P* < 0.05; **Fig. 5C**). Similarly, the effect size of sex also increased in the treated groups. Specifically, among the 18 NER genes that showed higher expression in the untreated ovary (Cohen’s d > 0), this effect was amplified for 16 genes in the PBS group and for all 18 genes in the MNU and ENU groups (**Fig. 5D** and **table S5**). Consistently, the ratios of effect sizes were significantly higher in the MNU and ENU groups compared to the PBS group (*t*-test based on log2-transformed values, both *P* < 0.001). Although baseline NER activity appears comparable between sexes, females still exhibit a stronger transcriptional response to exogenous pressure, potentially contributing to more efficient repair. Since BER and NER are essential for repairing alkylation-induced lesions, their low activity in males provides a compelling explanation for the elevated mutation rates observed after MNU and ENU treatment.

Above we performed bulk RNA sequencing on reproductive organs that contains both germ cells and supportive somatic cells (some of which provide proteins and transcripts to germ cells). To further discern whether the observed disparity in gene expression was intrinsic to the germ cells, we leveraged public single-cell RNA sequencing datasets of untreated female and male germ cells in mice (*60–62*). Consistent with the above findings, we found that the BER and NER genes are both expressed at significantly higher levels in the female germ cells than the male germ cells (Paired *t*-test, both *P* < 0.001; **fig. S5** and **table S6**). These findings suggest that the transcriptional differences of DNA repair pathways are an inherent property of germ cells, in line with the compromised repair capacity and increased susceptibility to mutagens in the male germline observed above.

## Discussion

The male-biased germline mutation rate is a well-documented phenomenon across vertebrates, yet its causes, whilst conventionally thought to have been resolved, more recently have been rendered uncertain. Our study provides experimental evidence supporting the damage-induced hypothesis (*1*, *33*, *39*), which posits that the male germline is more susceptible to damage compared to the female germline. Indeed, while our experimental procedures ensured equal systemic administration of mutagens between female and male, as females are a little smaller the actual stress experienced by ovary should, all else being equal, have been a little higher, rendering the higher mutation rate in males a conservative result. This is consistent with either differential repair of lesions (less common in males) or differential exposure (more common in males), possibly owing to anatomical differences between testes and ovaries. Our data argue against the latter differential exposure argument and for the former differential repair model as we see increased expression of repair genes in ovary than in testes, while the mutation rates show the reverse tendency. Indeed, the ovary exhibited significantly greater expression changes than the testis (*t*-test, *P* < 0.001; **fig. S6**), even under non-mutagenic PBS treatment, suggesting that females may experience stronger stress. The more parsimonious model then is that both germlines “see” the mutagens but female germline responds more to repair damage.

Distinguishing damage-induced mutations from replication errors remains challenging, not least because BER enzymes such as NEIL1 repair oxidized bases in the replicating template and are part of the replication complex (*63*). Nonetheless, multiple lines of evidence point to damage induction as the primary driver of elevated mutation rates in the male germline. In MNU- and ENU-treated males showing significantly higher germline mutation rates, we identified specific enrichment of two mutational signatures (SBS12 in MT-MNU and SBS30 in MT-ENU). While SBS30 is an established marker of defective DNA damage repair (*64*, *65*), SBS12 also appears to originate from damage-induced processes rather than replication errors. Crucially, SBS12 lacks the replicational strand asymmetry characteristic of replication errors but instead exhibits pronounced transcriptional strand asymmetry, suggesting involvement of transcription-coupled repair mechanisms (*66*). Furthermore, the signature’s predominant occurrence in post-mitotic liver cancers argues strongly against a replication-dependent origin (*66*). Supporting these observations, our transcriptome analysis, both bulk and single cell, revealed consistently lower expression of DNA repair genes in testes compared to ovaries, particularly in the BER pathway. This inherent deficiency became more pronounced following mutagen exposure, demonstrating male germ cells’ reduced capacity to cope with DNA damage. On exposure, the NER pathway also showed an increased sex bias. Since BER and NER are essential for repairing alkylation-induced lesions, their impaired activity in males provides a parsimonious explanation for the elevated mutation rates observed after MNU and ENU treatment.

Given the complexity and variability of both endogenous and exogenous mutagens experienced by natural populations, the mutagenic chemicals used in this study do not fully replicate the conditions encountered in nature. Nevertheless, our study introduces exogenous mutagenic stress in a well-controlled setting, allowing our experimental conditions to serve as an exemplar for uncovering potential sex differences in DNA repair capacity. Moreover, the DNA alkylation adducts caused by MNU/ENU are similar to certain types of damage from endogenous sources (*67*). The fact that DNA repair genes from BER pathway are expressed at lower levels in the testes of untreated mice further supports the broader relevance of our findings to the male-biased mutation patterns observed in nature.

The evidence that differential repair rates are seen between the sexes in germline raises a further evolutionary problem, namely why this might be. Previously, when it was considered that the male bias was owing solely to mutagenic effects of replication, the male bias was implicitly seen as an undesirable but unavoidable consequence of the highly mitotic nature of sperm production compared with oocyte production (limited to few mitoses between zygote and oocyte). Similarly, if the male germ line is more exposed to mutagens than the female germline the male female mutation rate difference could then also be configured an undesirable but possibly unavoidable consequence of a difference in male/female anatomy. Indeed, protamine packaging of DNA in the sperm head enables DNA compaction (and protection against damage (*68*)) but also an inability to perform repair. Failure of DNA protamination can then leave DNA vulnerable to damage (*69*) that requires BER correction after fertilization. Likewise, apoptosis to removes more heavily damaged cells (*70*) may fail and let through too many, also giving a male bias. Indeed, in humans there is a truncated sperm BER pathway relying on oocyte mediated repair (*71*) to complete the process.

While differences in sperm and oocyte maturation may explain some increased mutagenicity in male germ line, the different levels of repair enzymes question why males don’t invest as much into germline repair as females do? There seems nothing obviously unavoidable about not expressing repair enzymes, at least during early stages of spermatogenesis. Indeed, a Chinese remedy for male infertility upregulates the BER pathway (*72*), indicating that higher expression levels are possible. While assumptions about the causes of the generation time effect (*3*) and the expected relationship between life history and mutation rate (*10–12*) require reevaluation , the male mutation bias problem now poses a new paradox: why do males have low levels of BER repair enzymes and why don’t mutagens cause upregulation similar to that seen with the Chinese remedy?

## Methods

### Mutagenic treatment

The experiment was conducted using wild-type C57BL/6J mice purchased from the Qizhen Technology (Hangzhou). Animals were kept under standard breeding conditions, with controlled temperature and lighting, and were provided with food and water *ad libitum*. At four weeks of age, both male and female mice were treated with mutagens (dosage specified in Fig. 1B) or PBS (0.1 mL per dose) via intraperitoneal injection. Each treatment group consisted of five males and five females, treated five days a week for three weeks. The mice were weighed at seven weeks of age following the treatments. Male mice consistently weighed more than female mice across all treatment groups, with an average male-to-female weight ratio of 1.16. Given male mice were heavier than females, but both sexes received the same absolute dose, the effective dosage per unit of body weight was slightly higher in females. This makes our conclusion that mutation rates are higher in males under certain treatments more conservative. Following one-week rest, each treated mouse at eight-week of age was mated with an untreated mouse of the same age to obtain offspring. The protocols of mouse care and experiments were approved by the ethics committee of the Deruikang institute (Reference Number: DRK2023150286).

### Sample collection and whole-genome sequencing

Muscle tissue specimens were collected from the offspring and their corresponding mating pairs. Genomic DNA was extracted using the HiPure Universal DNA Kit (Magen, China) following the manufacturer’s instructions. Libraries were prepared using Covaris LE220R-plus (Covaris, USA) and AMpure XP system (Beckman Coulter, USA). Whole-genome paired-end sequencing was performed on the Illumina PE150 platform (Astrocyte Technology, Hangzhou) to generate 150 base-pair paired-end reads. Each sample was sequenced to achieve a target coverage of 40×. A total of 51 mating pairs and 91 of their offspring were sequenced and retained in the final dataset (**table S1**). The sequencing reads have been deposited in a public database and will be made available upon acceptance of the manuscript.

### Variant calling and filtering

Sequencing reads from each sample were aligned to the mouse reference genome (GRCm39) using the BWA-MEM algorithm version 0.7.17-r1188 (*73*). The aligned reads were sorted using SAMtools version 1.13 (*74*). The duplicated reads as well as reads mapped to multiple regions of the genome were removed using Picard MarkDuplicates version 3.1.1 (https://broadinstitute.github.io/picard/). Aligned reads were filtered using SAMtools view with the parameter “-b -h -q 20 -f 3 -F 4 -F 8 -F 256 -F 1024 -F 2048”. Variants for each trio (i.e. treated parent sample, untreated parent sample and one sample of their offspring) were called from HaplotypeCaller with GATK version 4.1.0 (*75*). The mutations identified in the offspring were filtered as followed:

(1) The variants were first filtered using a QUAL score threshold of > 200.
(2) Only sites in which both parents were homozygous for the reference allele, and the offspring was heterozygous were kept, to select variants that conformed to Mendelian inheritance.
(3) Only positions with a DP > 20 for each individual in the trio were kept, to minimise the effects of sequencing errors in regions of low sequencing depth.
(4) Only variants in which the offspring had 30–70% of the reads supporting the alternative allele were kept.
(5) Only variants in which the mutant reads were not detected in either parental sample were kept.
(6) Only variants in which the offspring had at least three reads supporting both the reference and alternative allelic on both forward and reverse strands were kept.
(7) Only variants located on autosomes were kept.

These quality control steps yielded reliable results, which were verified by manual inspection using SAMtools tview. To further confirm the reliability of the mutations, seven randomly selected mutations were validated by Sanger sequencing. A mutation was considered verified if both parents exhibited the reference genotype and the offspring exhibited the heterozygous mutant genotype. All seven mutations were successfully verified. A comprehensive list of the mutations identified in this study is provided in table S2.

### Analysis of mutation rate and spectrum

To estimate SNV rates per site, we calculated the number of SNV per callable genome (defined as sites with a depth > 20, multiplied by two for diploid genomes). To access the mutagen-induced mutation rate, the mutation rate of each sample was adjusted by subtracting the mean mutation rate of the sex-matched PBS-treated trios, yielding the mutation rate relative to PBS.

We applied the Poisson test to assess the significance of differences in SNV rates between males and females under the same treatment, as well as between different mutagens and PBS within the same sex, obtaining p-values. To compare SNV rates relative to PBS, we used the Wilcoxon rank-sum test. All p-values were adjusted using the Benjamini-Hochberg (BH) method to control the false discovery rate.

To investigate mutation spectrum, we extracted six types of SNVs from each group and quantified the occurrence of each SNV type, using Chi-square tests to assess differences in SNV profiles between mutagen-treated and PBS-treated trios within the same sex, as well as between female-treated and male-treated trios under the same treatment.

For the SBS signature analysis, we performed the following analyses using R (version 4.4.1) and the sigfit package (version 2.2). We first visualized 96 types of single base substitutions (SBS) in both FT and MT under PBS, MNU, and ENU treatments using the plot_spectrum function. We then applied the fit_signatures function which employs Markov Chain Monte Carlo (MCMC) sampling to fit a set of predefined mutational signatures to the observed mutational catalogues, to infer the potential mutational processes contributing to the observed spectra.

### Transcriptomic analysis of ovary and testis

To further validate the observed sex-specific differences in DNA damage repair capabilities in MNU/ENU-treated trios and to decipher the potential underlying mechanisms at the transcriptome level, C57BL/6J mice were exposed to PBS, MNU, and ENU following the same treatment protocols described above. Each treatment group included six males and six females, started at four weeks of age. Mice received intraperitoneal injections for two weeks (five days per week), followed by gonad sampling. For untreated controls, gonads were collected from age-matched, untreated mice (six males and six females).

Total RNA was extracted from the sampled tissues using Trizol reagent Kit (Invitrogen, USA) following the manufacturer’s instructions. RNA-seq libraries were prepared using NEBNext Ultra RNA Library Prep Kit for Illumina (New England Biolabs, USA), and sequencing was performed on the Illumina Novaseq 6000 platform (Astrocyte Technology, Hangzhou). On average, 32.1 million paired reads were obtained per sample. The sequencing reads have been deposited in a public database and will be made available upon acceptance of the manuscript.

Sequencing reads were aligned to the mouse reference genome (GRCm39) with STAR version (*76*) using Ensembl gene annotations. Gene-level read counts were generated using featureCounts version (*77*) and were subsequently normalized to transcripts per million base pair (TPM).

To investigate the capacity for DNA protection and repair in the germline, we adopted a curated set of human DNA repair (n = 219) as reported (*56*) and described in the updated version (https://www.mdanderson.org/documents/Labs/Wood-Laboratory/human-dna-repair-genes.html).

These genes were then converted to their mouse orthologs via the Ensembl BioMart database using RefSeq mRNA IDs, yielding a list of 214 mouse genes. TPM values for these genes were extracted from the normalized expression matrix.

Expression patterns across sexes and treatments were visualized using the pheatmap R package (*78*). For each gene under each treatment, Cohen’s d was calculated between males and females using the Effsize R package (*79*), and the corresponding *P* values were obtained via t-tests.

## Code availability

All code is available at https://doi.org/10.6084/m9.figshare.29553662.v1

## Data availability

Sequence generated for this project is deposited at the China National Genomics Data Center under the project number PRJCA039748.

## Acknowledgements

Haoxuan Liu is supported by National Key Research and Development Program of China (2024YFA1802500) and National Natural Science Foundation of China (32470646).

## Author contributions

Haoxuan Liu, Laurence D. Hurst, and Danqi Qin conceived the project and designed the experiment. Danqi Qin carried out the experiment and analyzed the data. Danqi Qin, Haoxuan Liu, and Laurence D. Hurst wrote the manuscript. All authors have read and approved the manuscript.

**The authors declare no conflict of interest.**

